# Cross-Individual Affective Detection Using EEG Signals with Audio-Visual Embedding

**DOI:** 10.1101/2021.08.06.455362

**Authors:** Zhen Liang, Xihao Zhang, Rushuang Zhou, Li Zhang, Linling Li, Gan Huang, Zhiguo Zhang

## Abstract

EEG signals have been successfully used in affective detection applications, which could directly capture brain dynamics and reflect emotional changes at a high temporal resolution. However, the generalized ability of the model across individuals has not been thoroughly developed yet. An involvement of other data modality, such as audio-visual information which are usually used for emotion eliciting, could be beneficial to estimate intrinsic emotions in video content and solve the individual differences problem. In this paper, we propose a novel deep affective detection model, named as EEG with audio-visual embedding (EEG-AVE), for cross-individual affective detection. Here, EEG signals are exploited to identify the individualized emotional patterns and contribute the **individual preferences** in affective detection; while audio-visual information is leveraged to estimate the **intrinsic emotions** involved in the video content and enhance the reliability of the affective detection performance. Specifically, EEG-AVE is composed of two parts. For EEG-based individual preferences prediction, a multi-scale domain adversarial neural network is developed to explore the shared dynamic, informative, and domain-invariant EEG features across individuals. For video-based intrinsic emotions estimation, a deep audio-visual feature based hypergraph clustering method is proposed to examine the latent relationship between semantic audio-visual features and emotions. Through an embedding model, both estimated individual preferences and intrinsic emotions are incorporated with shared weights and further are used together to contribute to affective detection across individuals. We conduct cross-individual affective detection experiments on two well-known emotional databases for model evaluation and comparison. The results show our proposed EEG-AVE model achieves a better performance under a leave-one-individual-out cross-validation individual-independent evaluation protocol. EEG-AVE is demonstrated as an effective model with good generalizability, which makes it a power tool for cross-individual emotion detection in real-life applications.

## I. Introduction

Electroencephalography (EEG) provides a nature way to record human brain activities and has been widely used in the affective intelligence studies [1]–[5]. In recent years, deep neural network learning methods have provided an effective and efficient approach to characterize informative deep features from EEG data and have achieved promising results in EEG-based affective detection applications. For example, a novel dynamic graph convolutional neural network (DGCNN) was proposed in [1] to learn the discriminant and hidden EEG characteristics in a non-linear approach for solving the multi-channel EEG based emotion decoding problem. Jirayucharoensak *et al*. [6] adopted a stack of several autoencoder structures to perform EEG-based emotion decoding and showed the deep learning network outperformed the traditional classification models such as support vector machine (SVM) and naïıve Bayes classifiers. The valid, useful and optimal EEG information can be explored in a deep belief network (DBN) structure, which was demonstrated to be beneficial to the decoding performance [7]. Cui *et al*. [8] proposed an end-to-end regional-asymmetric convolutional neural network (RACNN) to capture the discriminant EEG features covering temporal, regional, and asymmetric information. Based on a series of pretrained state-of-the-art CNN architectures, Cimtay and Ekmekcioglu [4] improved the feature extraction performance and classification capability based on raw EEG signals. The existing literature has shown deep learning is a powerful tool in EEG processing, which captures the abstract representations and disentangle the semantic gap between EEG signals and emotion states.

However, due to the problem of individual differences, the stability and generalizability of EEG-based affective detection models are of great challenge. Especially, EEG data are very weak signals and easily susceptible to interference from undesired noises, making it different to distinguish individual-specific and meaningful EEG patterns from noise. The key to solving the problem of individual differences is to minimize the discrepancy in feature distributions across individuals. To improve model generalization to the variance of individual characteristics, transfer learning methods have been introduced and a fruitful line of prior studies has been explored [2], [9]–[11]. Based on feature distribution and classifier parameters learning, Zheng and Lu [10] developed two types of subject-to-subject transfer learning approaches and showed a significant increase in emotion recognition accuracy (conventional generic classifier: 56.73%; the proposed model: 76.31%). Lin and Jung [11] proposed a conditional transfer learning framework to boost a positive transfer for each individual, where the individual transferability was evaluated and effective data from other subjects were leveraged. Li *et al*. [2] developed a multi-source transfer learning method, where two sessions (calibration and subsequent) were involved and the data differences were transformed by the style transfer mapping and integrated classifier. Among various transfer learning strategies, domain adaptation is a popular way to learn common feature representations and make the feature representations invariant across different domains (source and target domains). Ganin *et al*. [12] proposed an effective domain-adversarial neural network (DANN) to align the feature distributions between source domain and target domain and also maintain the information of the aligned discriminant features which are predictive of the labels of source samples. Instead of the conventional domain adaptation methods that adapted a well-trained model based on a specific domain to another domain, DANN could well learn the shareable features from different domains and maintain the common knowledge about the given task. Inspired by this work, Li *et al*. [13] proposed a bi-hemisphere domain adversarial neural network (BiDANN) model for emotion recognition using EEG signals, in which a global and two local domain discriminators worked adversarially with an emotion classifier to improve the model generalizability. Li *et al*. [3] proposed a domain adaptation method through simultaneously adapting marginal and conditional distributions based on the latent representations and demonstrated an improvement of the model generalizability across subjects and sessions.

On the other hand, with the great development and application of the internet and multimedia nowadays, there are many approaches to characterize audio-visual content and embed the conveying information with other feature modalities for emotion detection. For example, based on traditional handcrafted audio and visual features, Wang *et al*. [14] investigated several kernel based methods to analyze and fuse audio-visual features for bimodal emotion recognition. Mo *et al*. [15] proposed Hilbert-Huang Transform (HHT) based visual and audio features for a time-frequency-energy description of videos and introduced cross-correlation features to indicate the dependencies between the visual and audio signals. Furthermore, the recent success of deep learning methods in computer vision brings new insights into video-content based affective study. Acar *et al*. [16] utilized CNNs to learn mid-level audio-visual feature representations for affective analysis of music video clips. Zhang *et al*. [17] proposed a hybrid deep model to characterize a joint audio-visual feature representation for emotion recognition, where CNN, 3D-CNN, and DBN were integrated with a two-stage learning strategy.

In general, current affective computing models can be mainly categorized into two streams. One stream is to predict individual preferences through analyzing a user’s spontaneous physiological responses (i.e. EEG signals) while watching the videos [18], [19]. The individualized reactions to emotions are well-considered, and an assumption is made here that different emotions could be elicited for different viewers when watching the same video. However, spontaneous response-based individual preferences prediction would be sensitive to individual differences and fail to achieve reliable performance in affective detection across individuals. Another stream is to estimate intrinsic emotions from video content itself by integrating visual and audio features in either feature-level fusion or decision fusion and building a classifier for distinguishing emotions [17], [20]. The video content-based intrinsic emotions estimation could achieve a stable emotion detection performance, but it fails to consider the deviations of individuals in emotion perceiving. This motivates us to study the underlying associations among emotions, video content, and brain responses, where video content functions as a stimulation clue indicating what kind of emotions would possibly be elicited and brain responses reveal individual emotion perceiving process showing how we exactly feel the emotions. An appropriate embedding strategy of individual preferences and intrinsic emotions in cross-individual affection detection tasks could be helpful to learn reliable affective features from video content and benefit to enhancing the estimation stability of individual emotions.

Besides, compared to unimodal analysis, multimodal fusion could provide more details, compensate for the incomplete information from another modality, and develop advanced intelligent affective systems [21]. Recently, Wang *et al*. [22] incorporated video information and EEG signals to improve the video emotion tagging performance. This study characterized a set of traditional visual and audio features, including brightness, color energy and visual excitement for visual features, and average energy, average loudness, spectrum flow, zero-crossing rate (ZCR), standard deviation of ZCR, 13 Mel-Frequency Cepstral Coefficients (MFCC) and the corresponding standard deviations for audio features. The proposed hybrid emotion tagging approach was realized on a modified SVM classifier, and the corresponding performance was improved from 54.80% to 75.20% for valence and from 65.10% to 85.00% for arousal after a fusion of multi-modality data. Inspired by the success of the embedding protocol across different data modalities, this study proposes a novel affective information detection model (termed as EEG-AVE) to learn transferable features from EEG signals **individual preferences prediction** with an embedding of affective-related multimedia characteristics (**intrinsic emotions estimation**) to enhance cross-individual affective detection performance. The proposed EEG-AVE model is illustrated in Fig. 1, which is composed of three parts: EEG-based individual preferences prediction, audio-visual based intrinsic emotions estimation, and multimodal embedding. **(1) EEG-based individual preferences prediction:** In this part, we propose a multi-scale domain adversarial neural network (termed as MsDANN hereinafter) based on DANN [12] to enhance the generalization ability of EEG feature representation across individuals and boost the model performance on individual preferences prediction. Specifically, EEG data from different individuals are treated as domains, where the source domain refers to the existing individuals and the target domain refers to the newcoming individual(s). Based on the input multi-scale feature representation, the feature extractor network, task classification network, and discriminator network are designed to make the source and target domains share similar and close latent distribution to work with the same prediction model. As the mining of emotional informative and sensitive features from EEG signals is still a great challenge, this study introduces a multi-scale feature representation to improve feature efficacy and model adaptability to complex and dynamic emotion cases. Compared to single-scale feature representation, pioneer studies have shown EEG signals analysis with such a coarse-grain procedure could be beneficial to emotion studies [23]–[25]. **(2) Audio-visual based intrinsic emotions estimation:** To enhance the model stability in cross-individual affective detection tasks, audio-visual content analysis is conducted to digest the intrinsic emotion information involved in the videos which could be used as supplementary information for individual affective detection. Due to the well-known “semantic gap” or “emotional gap” that the traditional handcrafted features may fail to sufficiently discriminate emotions, we develop a deep audio-visual feature based hypergraph clustering method (termed as DAVFHC) for characterizing semantic and high-level audio-visual features. Here, two pretrained CNN architectures (VGGNet [26] and VGGish [27], whose performance have been widely recognized in audio-visual information analysis [28], [29]) are adopted to explore the emotion-related audio-visual characteristics and the most optimal features are fused through a hypergraph theory. **(3) Multimodal embedding:** The final affective detection result is determined by an embedding model where the predicted individual preferences from EEG signals and the estimated intrinsic emotions from audio-visual content are fused at a decision level. The compensation information from different modalities contributes together to tackle the individual differences problems in affective detection.

**Fig. 1:**
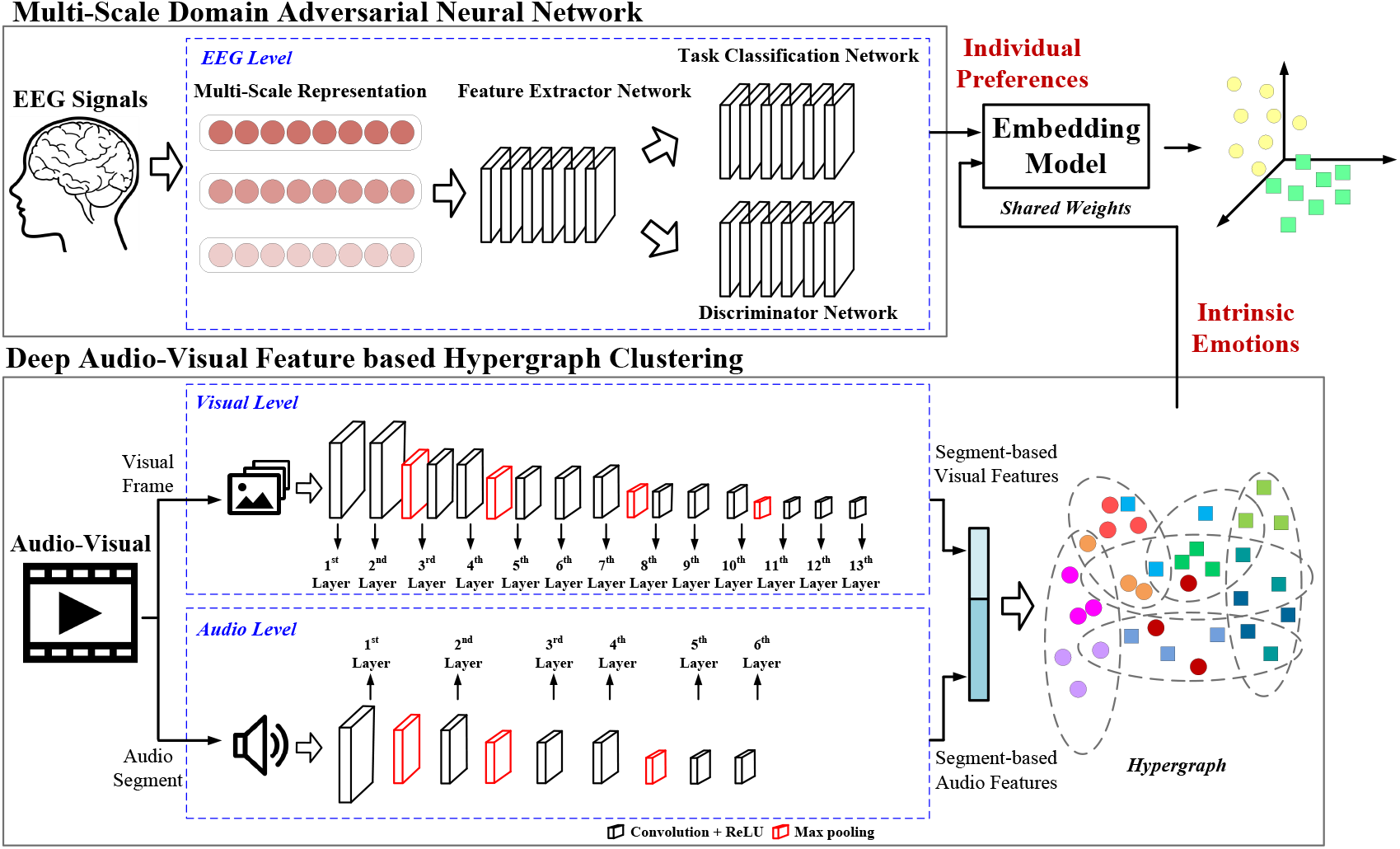
The proposed EEG-AVE model.

The major contributions of this work are summarized as follows. (1) We propose a novel cross-individual affective detection model (EEG-AVE) to incorporate spontaneous brain responses and stimulation clues in a hybrid embedding strategy. Both EEG and audio-visual information are exploited to digest different dimensions of emotions, and the compensation relationships among different data modalities on the affective detection study are examined. (2) We introduce an effective individual preferences prediction method (MsDANN) to estimate individual emotions from EEG signals, where the impact of individual differences is diminished through a transfer learning approach. (3) We present an efficient intrinsic emotions estimation method (DAVFHC) to characterize the emotion-related in audio-visual materials as the supplementary information for the cross-individual affective study. Here, the semantic audio-visual features are extracted by using deep learning methods, and the complex and latent relationships of deep audio-visual features with emotion labels are measured with hypergraph theory.

## II. Methodology

## A. Individual Preferences Prediction

In this section, we propose a new transfer learning based neural network, MsDANN, to address the individual differences problem in EEG based emotion detection. In this network, a multi-scale feature representation is incorporated to capture a series of rich feature characteristics of EEG signals and maximize the informative context for predicting a diverse set of individual preferences in emotions. Specifically, we extract the differential entropy (DE) features [30] from the defined frequency sub-bands (refer to Table I) at different frequency/scale resolutions (1 Hz, 0.5 Hz, and 0.25 Hz), and build respective domain adaptation models with domain adversarial training methods. In the proposed MsDANN, the common features from different individuals are learnt; at the same time, the relationships between the learnt common features and the related emotion information are preserved. The network structure of MsDANN is shown in Fig. 2, which is composed of three parts: the generator (feature extractor network) for deep feature extraction, the classifier (task classification network) for emotion label prediction, and the discriminator (discriminator network) for real or fake data distinguishing. Here, the generator and classifier could be considered as a standard feed-forward architecture, while the generator and discriminator are trained based on a gradient reversal layer to ensure the feature distributions of two domains as indistinguishable as possible. In this study, the EEG data with emotion labels are treated as the source domain to train the generator, classifier and discriminator; while the EEG data without emotion labels are utilized to train the generator and discriminator. Through this multi-scale deep framework, a set of transferable features involving affective information could be characterized, the cross-domain discrepancy could be bridged, and the classification performance could be effectively improved in both source and target domains.

**TABLE I:** The defined frequency sub-bands for DE feature characterization.

| | $\theta$ | $\alpha 1$ | $\alpha 2$ | $\beta 1$ | $\beta 2$ | $\beta 3$ | $\gamma 1$ | $\gamma 2$ | $\gamma 3$ |
| --- | --- | --- | --- | --- | --- | --- | --- | --- | --- |
| frequency band (Hz) | 4-8 | 8-10 | 10-13 | 13-16 | 16-20 | 20-28 | 28-34 | 34-39 | 39-45 |

**Fig. 2:**
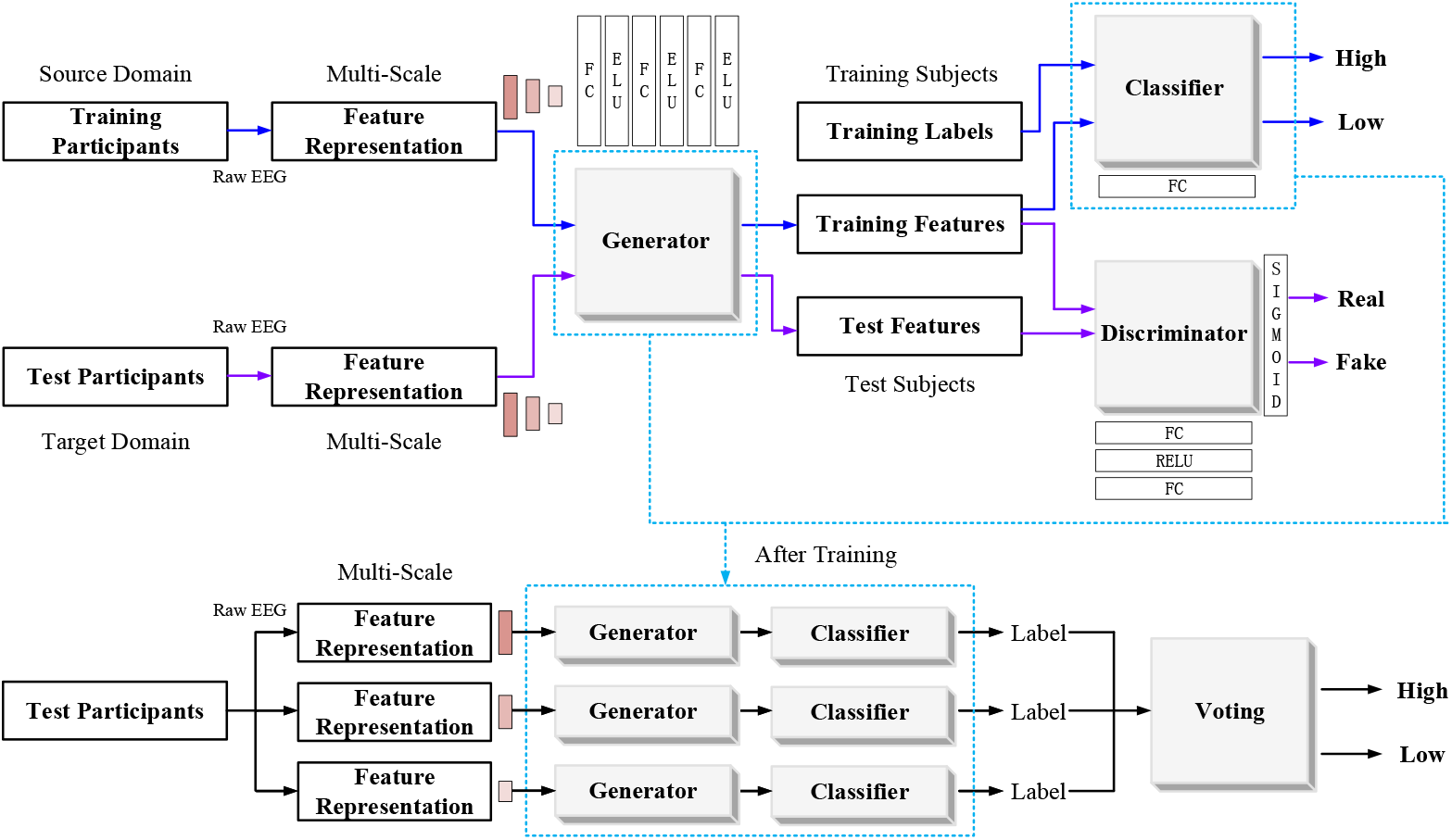
The proposed MsDANN model.

To learn a shared common feature space between the source and target domains and also guarantee the learnt feature representation involving enough information for revealing the emotion states, the loss objective function is designed below. Suppose that the source and target domains are denoted as 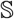 and 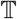. In the domain learning, the EEG data with emotion labels in 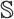 are given as 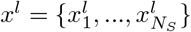 and 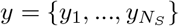, where 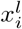 is the input EEG data at *l*th scale feature representation and *y_i_* is the corresponding emotion label of 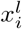. *N_S_* is the sample size of *x^l^*. On the other hand, the unlabeled EEG data in 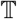 is denoted as 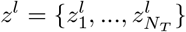, where 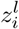 is the input EEG data at *l*th scale feature representation and *N_T_* is the corresponding sample size of *z^l^*. We denote the generator, classifier, and discriminator as *r_θ_*, *c_σ_*, *d_µ_* with the parameters of *θ*, *σ* and *µ*. To ensure the learnt features by *r_θ_* from source domain or target domain are indistinguishable, the domain adversarial training objective function is given as

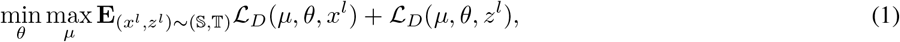

where 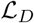 is a binary cross-entropy loss for the discriminator to be trained to distinguish 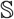 and 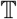, defined as

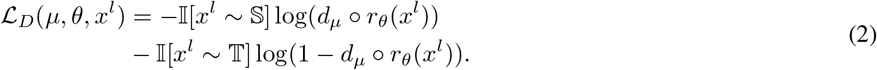

Here, 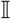 is an indicator function. Based on Eq. 1, we add another loss function 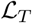 for the classifier part as

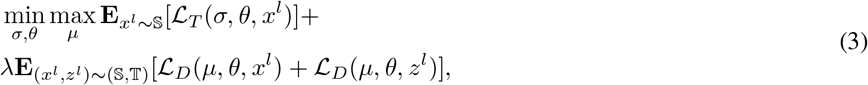

where 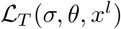 is the classification loss in the source domain, determined by parameter during the learning process, given as Σ*Loss*(*c_σ_* ◦ *r_θ_*(*x^l^*), *y*). *λ* is a balance parameter during the learning process, given as

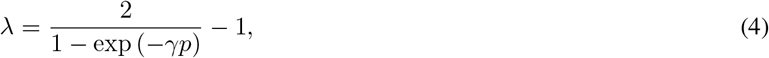

where *γ* is a constant value and *p* is a factor of epoch. Eq. 3 is the final objective function for MsDANN model training. The proposed MsDANN model is an end-to-end framework for cross-individual emotion prediction based on EEG signals, combining the feature learning adaptation and emotion classification into a unified deep model. Based on the input data with multi-scale DE feature representation, the domain adaption and classification loss are exploited to guide the generator to learn effective feature representations across individuals via the gradient reversal layer and efficiently tackle the individual differences problem in EEG data processing.

## B. Intrinsic Emotions Estimation

At present, a number of well trained deep CNN models have been successfully applied to multimedia processing, such as AlexNet [31], GoogLeNet [32] and VGG [26] for visual content, and VGGish [27] for audio content. The deep features could bridge the semantic gap and improve semantic interpretation performance. In this section, we develop a DAVFHC method to learn and decode the semantic features from audio-visual content for intrinsic emotions estimation.

At the visual level, a pretrained VGGNet network [26] is utilized to process frame-based visual information and characterize effective visual features. The training and testing data sets were based on ILSVRC-2012, with 1.3M training pictures, 50K test pictures, and 100K validation pictures. The network was trained by optimizing a polynomial logistic regression objective function with a smallest batch-based gradient descent momentum. Considering the balance of layer depth and performance, VGG16 is utilized in this paper to characterize the frame-based visual features. It consists of 13 convolutional layers and 3 fully connected layers. The corresponding number of convolution kernels at each layer are 64, 64, 128, 128, 256, 256, 256, 512, 512, 512, 512, 512, and 512, and the kernel size is 3 × 3. As illustrated in Fig. 3, the visual feature extraction procedure includes three steps. **1. Frame-based visual feature extraction.** The video frames are input to the pretrained VGG16 and the corresponding feature maps are characterized at each convolutional layer. For each layer, an average feature map is then calculated and converted into a feature vector. **2. Segment-based visual feature extraction.** Instead of direct averaging all the frame-based features in one segment, we introduce an adaptive key frame detection step to detect a key frame from every segment based on the feature distribution. Suppose that one segment is composed of *k* frames with the corresponding extracted features, denoted as 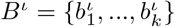, where *ι = 1, …, N_ι_* refers to the convolutional layer. The key frame detection is illustrated as follows. (1) All frames are grouped into one cluster in terms of *B^ι^*; (2) The cluster center *c^ι^* is computed; (3) The distance between each frame 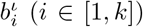 and the cluster center *c^ι^* is calculated, denoted as 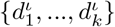; (4) the frame which is the closest to *c^ι^* is selected as the key frame of the segment, termed as 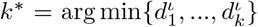. Then, the corresponding feature of the key frame 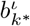 is treated as the segment-based feature representation. **3. Segment-based visual feature fusion.** The characterized segment-based features at each single convolutional layer 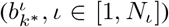 are then fused by concatenation. Empirically, the segment length is set to 1s. To get the semantic features, only the characterized features at the last two convolutional layers (*ι* = 12 and 13) are used as visual features (Ψ*_v_*) in the proposed DAVFHC method.

**Fig. 3:**
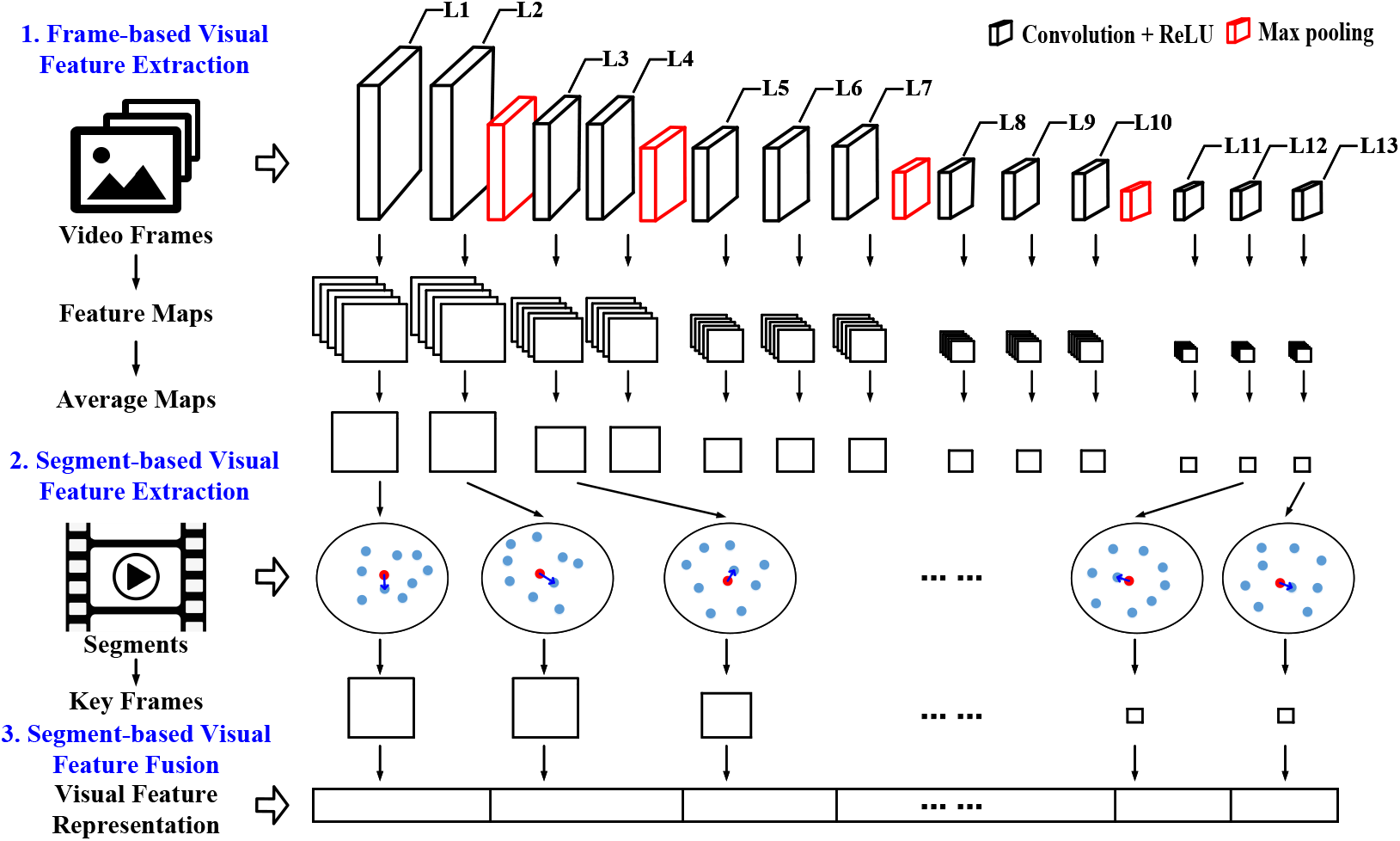
The visual feature extraction procedure.

At the audio level, a pretrained CNN network, VGGish [27], is adopted to characterize effective audio features. VGGish is a deep network model trained on a Youtube-8M database (training/validation/test: 70M/20M/10M), which has been proved to be capable of extracting effective and efficient deep auditory features in various applications [29], [33], [34]. The network contains 6 convolutional layers, and the corresponding numbers of convolution kernels are 64, 128, 256, 256, 512, and 512, respectively. The kernel size is 3 × 3. Same as the visual feature extraction process, the audio features are also characterized at the segment level. **1. Data preparation.** The audio signals are detected from the emotional clips and then partitioned into a number of segments with a fixed length. **2. Data preprocessing.** The segment-based audio data is preprocessed following the procedures presented in [27]. **3. Deep audio feature characterization.** For each segment, the logarithmic melspectrum is characterized and input to the VGGish. The deep feature maps are extracted at each convolutional layer and averaged into one feature map. **4. Deep audio feature fusion.** For each segment, the feature map at each single convolutional layer is converted into a feature vector. The converted feature vectors across different convolutional layers are then fused by concatenation. Empirically, the segment length is set to 1s (same as the visual data. To get the semantic features, only the feature vectors extracted from last two layers (5th and 6th) are used as audio features (Ψ*_A_*) in the proposed DAVFHC method.

The characterized segment-based visual and audio features are concatenated and formed into a segment-based audio-visual feature vector termed as Ψ*_M_ = [Ψ_V_,* Ψ*_A_*]. The complex relationships among all the segments from the emotional clips are constructed with a hypergraph which has been widely recognized as an effective approach for complex hidden data structure description. For the traditional graph, only pairwise relationships between any two vertices are considered, which would lead to the information loss [35]. In the hypergraph, one edge (termed as hyperedge in the hypergraph) could connect more than two vertices and the complex relationship among a group of vertices could be well described. In the paper, the segments are the vertices denoted as *V*, and the connections among the segments are the hyperedges denoted as *E*. One hypergraph could be represented as *G = (V, E*), where the vertices and hyperedges are denoted as *V = {v*_1_*, v*_2_*, …, v*_|*V* |_} and *E = {e*_1_, e_2_*, …, e*_|*E*|_}, respectively. The vertices belong to one hyperedge *e_k_* ∈ *E* is termed as 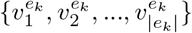. To define the vertices and hyerpedges relationships, the similarity between any two vertices (the emotional clip segments denoted as 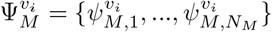 and 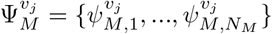, with the feature dimensionality of *N_M_*) are measured as

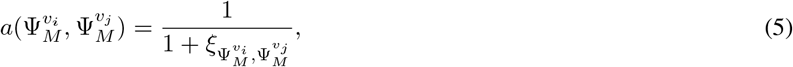

where 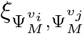 is the calculated distance, given as

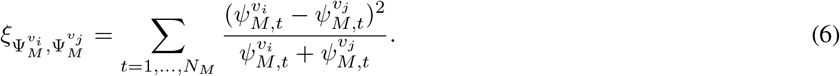

Based on the measured similarity matrix 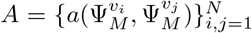 (*N* is the sample size), an incident matrix *H* ∈ |*V* | × |*E*| is formed, in which the connection relationships between the vertices *V* and the hyperedges *E* is described as

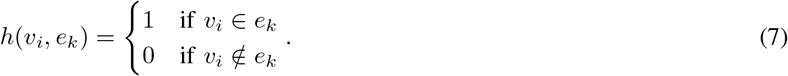

The hyperedge weight matrix *W* is a diagonal matrix indicating the weights of all the hyperedges *E* in the hypergraph *G*. The weight *w*(*e_k_*) of one hyperedge *e_k_ E* is computed based on the calculated similarities among the vertices that belong to *e_k_*, given as

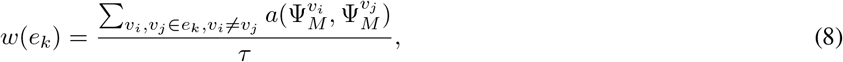

where 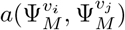 is the similarity value between the vertices of *v_i_* and *v_j_*, given in Eq. 5. *τ* is the total number of vertices connected to the hyperedge *e_k_*. As *w*(*e_k_*) is a measurement of all the similarity relationships among the vertices that belong to one hyperedge, a higher *w*(*e_k_*) value indicates a strong connection of homogeneous vertices of the hyperedge and a lower *w*(*e_k_*) refers to a weak connection of the hyperedge in which the connected vertices share little similar properties. In other words, the hypergraph structure could well describe the relationships of the audio-visual segments in terms of properties. The vertex degree matrix (*D_v_*) is a diagonal matrix presenting the degree of all the vertices in the hypergraph *G*. The degree of one vertex *v_k_ V* is calculated as the summation of all the hyperedge weights of the hyperedges that the vertex belong to, defined as

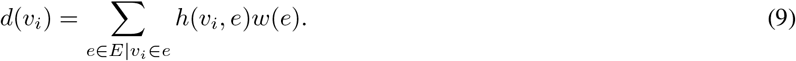

The hyperedge degree matrix (*D_e_*) is also a diagonal matrix showing the degree of all the hyperedges in the hypergraph *G*. The degree of one hyperedge *e_k_* ∈ *E* is calculated as the summation of all the vertices that connect to the hyperedge, given as

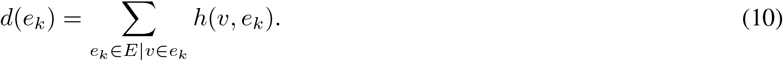

In this study, we introduce a spectral hypergraph partitioning method [36] to partition the constructed hypergraph into a number clusters corresponding to the emotion states (high or low). Thus, it is a two-way hypergraph partitioning problem which could be described as

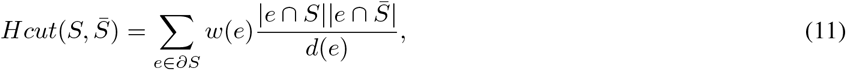

where *S* and 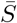 are the partitions of the vertices *V*. For two-way partitioning, 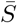 is the complement of *S*. *∂S* is the partition boundary, given as 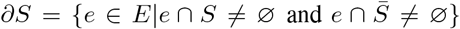. *d*(*e*) is the hyperedge degree defined in Eq. 10. To avoid unbalanced partitioning, 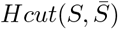 is further normalized by

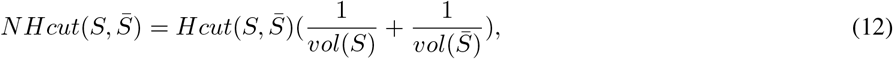

where *vol*(*S*) and 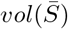 are the volumes of *S* and 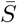, given as *vol*(*S*) = Σ_*v*∈S_ *d*(*v*) and 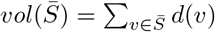. The partitioning rule is to look for the weakest hyperedge *e* between *S* and 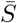, where the vertices in the same cluster should be tightly connected (high hyperedge weights) and the vertices in the different clusters should be weakly connected (low hyperedge weights). An optimal partitioning is given in Eq. 13 to find the weakest connection between two partitions, which is an NP-complete problem solved by a real-valued optimization method.

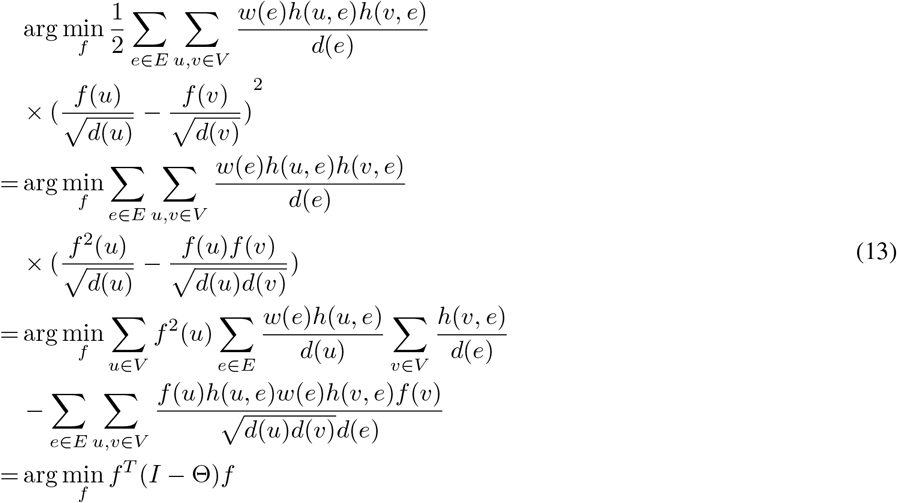

where Θ is given as

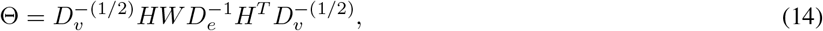

and *I* is an identity matrix with the same size as *W*. The hypergraph Laplacian is denoted as

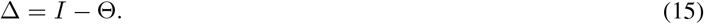

The optimal solution is transformed to find the eigenvectors of ∆ whose eigenvalues are the smallest. In other words, the optimal hypergraph partitioning results find the top eigenvectors with the smallest non-zeros eigenvalues in ∆ and form an eigenspace for the subsequent vertex clustering with the K-means method. Through this approach, all the vertices are grouped into two clusters. The corresponding emotion state of each cluster is determined by the majority distribution of the involved vertices. If most of vertices are belong to high level, the cluster’s emotion state is assigned as high; on the other hand, it is assigned as low. In practice, to avoid information leaking, the clusters’ emotion states are only determined based on the training samples.

### C. Embedding Model

Based on the aforementioned work, we incorporate the estimated intrinsic emotions based on deep audio-visual features and the predicted individual preferences from the collected simultaneous EEG signals, and conduct a decision-level information fusion for final affective prediction. Specifically, we fuse EEG signals and audio-visual information in a decision level through shared weights. Suppose that the predicted emotional individual preferences based on EEG signals are denoted as 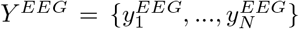 and the estimated intrinsic emotions based on audio-visual content are denoted as 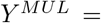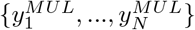 The final detected affective results are determined by

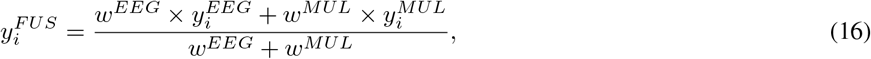

where *w^EEG^* and *w^MUL^* are the shared weights of EEG signals and audio-visual information in the fusion process. 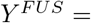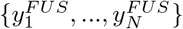 are the final affective detection results.

## III. Experimental Results

In this section, we conduct extensive experiments on MAHNOB-HCI [37] and DEAP [38] databases which are commonly used to evaluate the effectiveness of cross-individual affective studies. To cross-compare with other studies, two types of groundtruth data which are commonly used in the literature are adopted here to evaluate the experimental results. One is the **aggregated groundtruth**, where different participants watching one video are tagged with the same emotion label. Another is the **non-aggregated groundtruth**, where different participants watching one video are tagged with different emotion labels according to the corresponding subjective assessment. Different from the aggregated groundtruth, different participants would have different emotional feelings to the same video, due to the differences in background, experience, religion, education, and so on. In other words, the non-aggregated groundtruth could be more capable of reflecting the emotion dynamics in individuals and should be more encouraged to be used for affective detection evaluation.

### A. Emotional EEG Databases

The MAHNOB-HCI database [37] contains EEG data of 30 participants (male/female: 13/17; age: 26.06±4.39) from different cultural backgrounds. A total of 20 commercial film clips (duration: from 34.9s to 117s, with an average of 81.4s and a standard deviation of 22.5s) were selected for emotional eliciting. After the emotional clip playing, the participants were requested to give a subjective assessment about their emotions during watching the emotional clip using a score in the range of 1 to 9. During the experiment, EEG signals were simultaneously collected at a sampling rate of 256Hz, by using the Biosemi active II system with 32 Ag/AgCl electrodes placed according to the standard international 10-20 electrode system. Due to the data incompleteness of participants 3, 9, 12, 15, 16, and 26, only 24 participants are used in this paper.

The DEAP database [38] consists of 32 subjects’ EEG emotion data. A total of 40 music videos, with a fixed duration of the 60s, were selected for emotional eliciting. The corresponding subjective feedbacks on different emotion dimensions were collected for each music video. The EEG signals were recorded at a sampling rate of 512Hz from 32 active AgCl electrode sites according to the international 10-20 system placement.

### B. Experiment Protocols

To cross-compare with the results presented in the other studies, we utilize a fixed threshold of 5 for scores (in the range of 1 to 9) to discretize the subjective feedback into high and low levels (≥5 high; < 5 low) as the non-aggregated groudtruth. The aggregated groundtruth is an average of all the returned subjective feedback for one video. Two performance metrics, detection accuracy *P_acc_* and F1-Score *P_f_*, are used to validate the evaluation performance. *P_acc_* is an overall detection performance measurement and *P_f_* is a harmonic average of the precision and sensitivity which is less susceptible to the unbalanced classification problems. The corresponding definitions are given as

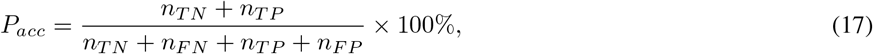

and

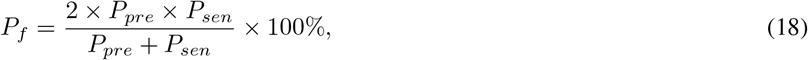

where *n_T N_* and *n_T P_* are the correctly predicted samples, and *n_F N_* and *n_F P_* are the incorrectly predicted samples. The precision *P_pre_* and sensitivity *P_sen_* are given as

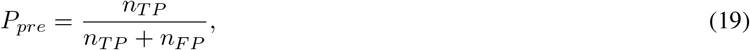

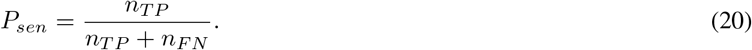

To fully evaluate the validity and reliability of the model performance, a strict leave-one-out cross-validation is adopted. All the predicted individual preferences and the estimated intrinsic emotions are obtained in a cross-validation manner. For the proposed MsDANN model, the model training and testing are conducted on a leave-one-individual-out cross-validation. In one round of cross-validation, all the samples from 1 individual are treated as the test data, while the other samples from the remaining individuals are used as the training data. Until each participant is treated as the test data once, the final result of MsDANN is a formation of all the obtained test results through the cross-validation rounds. For the developed DAVFHC method, the model training and testing are conducted on a strict leave-one-video-out cross-validation. In one round of cross-validation, all the samples from 1 video are used as test data and the other samples from the remaining videos are treated as training data. Until each video is treated as the test data once, the final prediction result of DAVFHC is a formation of the obtained test results in all the cross-validation rounds. In other words, after obtaining all the test results of all EEG and video samples in the above-mentioned cross-validation rounds, the final affective results are obtained by a decision fusion.

### C. Cross-Individual Affective Detection Experiments

To improve the affective detection performance, both EEG signals and audio-visual information are embedded in the proposed EEG-AVE model. Here, we roughly estimate what kind of emotion could be triggered according to the audio-visual content itself (intrinsic emotions estimation), and detect the individual preferences for each individual through analyzing the recording EEG signals while he / she is watching the multimedia material (individual preferences detection). The contributions of EEG signals and audio-visual information through the affective detection process are considered equally important. The corresponding emotion decoding performance for valence and arousal on MAHNOB-HCI and DEAP databases are reported in Table II. We compare EEG-AVE model with the existing representative methods such as [37], [39], [40], [41], and [22]. It is worth note that the experimental results presented in [22] were evaluated with the aggregated groundtruth.

**TABLE II:** Affective detection performance on MAHNOB-HCI and DEAP databases.

| Methods | Groundtruth | MAHNOB-HCI |  |  |  | DEAP |  |  |  |
| --- | --- | --- | --- | --- | --- | --- | --- | --- | --- |
|  |  | Valence |  | Arousal |  | Valence |  | Arousal |  |
| | | $P_{acc}$ | $P_f$ | $P_{acc}$ | $P_f$ | $P_{acc}$ | $P_f$ | $P_{acc}$ | $P_f$ |
| Soleymani *et al.* [37] | Non-Aggregated | 57.00 | 56.00 | 52.40 | 42.00 | - | - | - | - |
| Zhu *et al.* [39] | Non-Aggregated | 58.16 | 56.36 | 61.35 | 63.08 | - | - | - | - |
| Huang *et al.* [40] | Non-Aggregated | 62.13 | - | 61.80 | - | - | - | - | - |
| Rayatdoost and Soleymani [41] | Non-Aggregated | 71.25 | 62.08 | 61.46 | 50.60 | 59.22 | 56.68 | 55.70 | 50.02 |
| Wang *et al.* [22] | Aggregated | 75.20 | 73.80 | 85.00 | 82.40 | 71.10 | 68.60 | 79.00 | 69.20 |
| **Proposed EEG-AVE model** | Aggregated | **90.21** | **90.45** | **85.59** | **86.55** | **75.26** | **77.16** | 71.92 | **79.50** |
| **Proposed EEG-AVE model** | Non-Aggregated | 71.13 | **66.83** | **66.47** | **63.25** | **68.50** | **68.81** | 54.52 | **60.80** |

For the MAHNOB-HCI database, our proposed model outperforms the existing methods for valence, where the *P_acc_* and *P_f_* results are 90.21% and 90.45% for the aggregated groundtruth and 71.13% and 66.83% for non-aggregated groundtruth. For the results with non-aggregated groundtruth, even the obtained *P_acc_* values of our proposed EEG-AVE model and Rayatdoost and Soleymani [41]’s work are comparable, a better *P_f_* of our proposed EEG-AVE model is observed, where *P_f_* is 62.08% for Rayatdoost and Soleymani [41]’s work and 66.83% for our model (improved by 7.65%). For the results with aggregated groundtruth, our proposed EEG-AVE model increases the affective detection performance by 19.96% for *P_acc_* and 22.56% for *P_f_*, compared to Wang *et al*. [22]’s work. Similar promising emotion recognition performance is observed for arousal, where the *P_acc_* and *P_f_* results are 85.59% and 86.55% for the aggregated groundtruth and 66.47% and 63.25% for non-aggregated groundtruth. For aggregated groundtruth, the proposed EEG-AVE model performs better than Wang *et al*. [22], especially for F1-score (increased by 5%). For non-aggregated groundtruth, the EEG-AVE model also gains better performance than the existing methods on recognition accuracy. Besides, the above results show aggregated groundtruth leads to a higher detection performance compared to the non-aggregated groundtruth, as the individual differences in emotional feelings about the clip are not considered.

For the DEAP database, our proposed model outperforms the existing methods for valence in terms of both accuracy and F1-score. For aggregated groundtruth, the *P_acc_* and *P_f_* results are 75.26% and 77.16%; For non-aggregated groundtruth, the *P_acc_* and *P_f_* results are 68.50% and 68.81%. Even the affective detection accuracy for arousal is not as good as the existing methods, where *P_acc_* values are 71.92% and 54.52% for aggregated groundtruth and non-aggregated groundtruth, respectively. The obtained F1-score values are the highest, where *P_f_* values are 79.50% and 60.80% for aggregated groundtruth and non-aggregated groundtruth, respectively. Due to the imbalance data distribution of DEAP database [42], F1-score is a better and more important metric for classification models which can distinguish specific types of errors including false positives and false negatives.

## IV. Discussion and Conclusion

To fully study the EEG-AVE performance, we also compare the proposed model with different embedding strategies and domain adaption conditions. Besides, we also examine the effect of deep and handcrafted multimedia affective representations.

### A. Performance Evaluation of Embedding Strategy

We compare the affective detection performance when different embedding strategies are adopted. Here are three embedding strategies: EEG+Visual+Audio (the proposed EEG-AVE model), EEG+Visual (only visual information embedded with EEG signals), and EEG+Audio (only audio information embedded with EEG signals). The corresponding affective detection performances for valence and arousal with aggregated and non-aggregated groundtruth on MAHNOB-HCI and DEAP databases are summarized in Table III.

**TABLE III:**
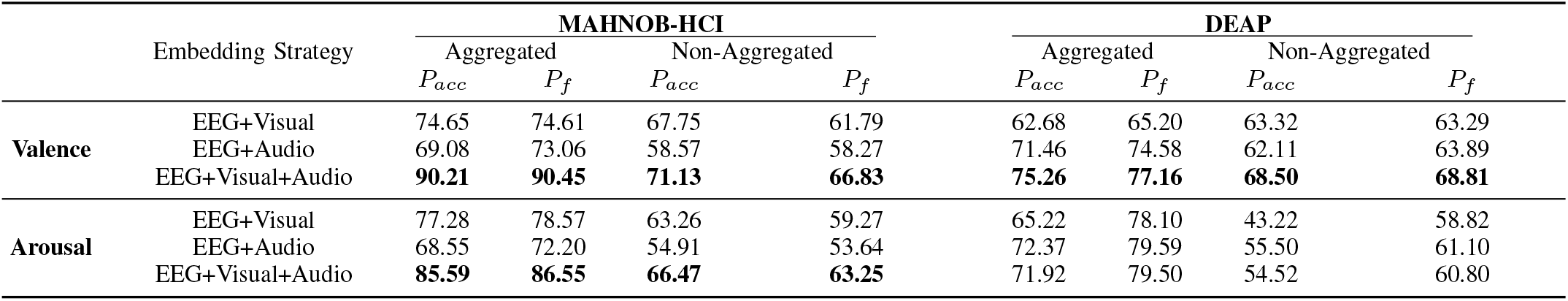
Affective detection performance with different embedding strategies on MAHNOB-HCI and DEAP databases.

The results on MAHNOB-HCI database show that EEG based affective detection with an embedding of both visual and audio information achieve the best performance for both valence and arousal. For EEG+Visual strategy, the affective detection performance for valence decreases to 74.65% (aggregated) and 67.75% (non-aggregated) for *P_acc_* and 74.61% (aggregated) and 61.79% (non-aggregated) for *P_f_*; while the affective detection performance for arousal decreases to 77.28% (aggregated) and 63.26% (non-aggregated) for *P_acc_* and 78.57% (aggregated) and 59.27% (non-aggregated) for *P_f_*. The average decrease rates of valence and arousal are 11.76% and 7.51%, respectively. For EEG+Audio embedding strategy, the affective detection performance for valence decreases from 90.21% to 69.08% for *P_acc_* and from 90.45% to 73.06% for *P_f_* when aggregated groundtruth is utilized; while it decreases from 71.13% to 58.57% for *P_acc_* and from 66.83% to 58.27% for *P_f_* when non-aggregated groundtruth is used. A similar decrease pattern is also observed on the affective detection performance for arousal, where it decreases from 85.59% to 68.55% for *P_acc_* and from 86.55% to 72.20% for *P_f_* when aggregated groundtruth is adopted; while it decreases from 66.47% to 54.91% for *P_acc_* and from 63.25% to 53.64% for *P_f_* when non-aggregated groundtruth is utilized. The average decrease rates of valence and arousal are 18.28% and 17.27%, respectively. The comparison results with different embedding strategies reveal that an embedding of both visual and audio information has a capability to reach better affective detection performance, compared to only visual or audio embedded. In addition, we find only visual embedded outperforms only audio embedded, which suggests that visual information plays a more critical role in emotion perceiving, especially in film clips.

For DEAP database, EEG-based affective detection with an embedding of both visual and audio information achieve the best performance for valence. For aggregated groundtruth, the *P_acc_* and *P_f_* values decrease from 75.26% and 77.16% (EEG+Visual+Audio) to 62.68% and 65.20% (EEG+Visual) and to 71.46% and 74.58% (EEG+Audio). The average decrease rate is 10.15%. For non-aggregated groundtruth, the *P_acc_* and *P_f_* values decrease from 68.50% and 68.81% (EEG+Visual+Audio) to 63.32% and 63.29% (EEG+Visual) and to 62.11% and 63.89% (EEG+Audio). The average decrease rate is 8.02%. However, for affective detection on arousal, similar results are obtained for EEG+Audio and EEG+Visual+Audio. One possible reason could be that the embedding strategies of visual and audio information could be different for different emotional dimensions. For example, it is observed that compared to visual information, audio plays a more important role for affective detection on the DEAP database, as the used stimuli for emotion evoking were music videos. For the MAHNOB-HCI database, the affection detection performance is more relied on visual information, as the used stimuli for emotion eliciting were movie clips.

### B. Performance Evaluation of Domain Adaptation Effect

To analyze the domain adaptation effect in solving the individual differences problem, we also introduce a baseline method, multi-scale neural network (termed as MsNN), for model comparison under the condition without deep domain adaption. Here, no feature adaption or transfer learning is adopted in EEG analysis, and the EEG based emotional individual preferences prediction is trained and tested on source domain and target domain separately. The corresponding affective detection performance of MsDANN and MsNN based EEG-AVE model for valence and arousal detection on MAHNOB-HCI and DEAP databases are reported in Table IV.

**TABLE IV:** Affective detection performance of MsDANN and MsNN on MAHNOB-HCI and DEAP databases using deep features.

|  | Embedding Strategy | EEG Model | MAHNOB-HCI |  |  |  | DEAP |  |  |  |
| --- | --- | --- | --- | --- | --- | --- | --- | --- | --- | --- |
|  |  |  | Aggregated |  | Non-Aggregated |  | Aggregated |  | Non-Aggregated |  |
| | | | $P_{acc}$ | $P_f$ | $P_{acc}$ | $P_f$ | $P_{acc}$ | $P_f$ | $P_{acc}$ | $P_f$ |
| **Valence** | EEG+Visual | MsDANN | 74.65 | 74.61 | 67.75 | 61.79 | 62.68 | 65.20 | 63.32 | 63.29 |
|  |  | MsNN | 70.04 | 73.03 | 60.98 | 59.20 | 58.21 | 66.07 | 52.15 | 58.69 |
|  | EEG+Audio | MsDANN | 69.08 | 73.06 | 58.57 | 58.27 | 71.46 | 74.58 | 62.11 | 63.89 |
|  |  | MsNN | 65.33 | 71.64 | 53.38 | 56.34 | 64.47 | 71.67 | 53.02 | 60.21 |
|  | EEG+Visual+Audio | MsDANN | 90.21 | 90.45 | 71.13 | 66.83 | 75.26 | 77.16 | 68.50 | 68.81 |
|  |  | MsNN | 82.00 | 83.95 | 63.58 | 62.35 | 66.03 | 72.55 | 55.59 | 61.84 |
| **Arousal** | EEG+Visual | MsDANN | 77.28 | 78.57 | 63.26 | 59.27 | 65.22 | 78.10 | 43.22 | 58.82 |
|  |  | MsNN | 72.02 | 75.76 | 55.14 | 54.93 | 65.20 | 78.11 | 42.45 | 58.30 |
|  | EEG+Audio | MsDANN | 68.55 | 72.20 | 54.91 | 53.64 | 72.37 | 79.59 | 55.50 | 61.10 |
|  |  | MsNN | 65.33 | 71.43 | 49.59 | 52.22 | 72.27 | 79.58 | 49.96 | 56.42 |
|  | EEG+Visual+Audio | MsDANN | 85.59 | 86.55 | 66.47 | 63.25 | 71.92 | 79.50 | 54.52 | 60.80 |
|  |  | MsNN | 78.39 | 81.42 | 58.00 | 58.17 | 72.62 | 80.07 | 50.62 | 57.59 |

For aggregated results on the MAHNOB-HCI database, compared to MsDANN based EEG-AVE model under EEG+Visual+Audio strategy, the detection performance of MsNN based EEG-AVE model decreases by 9.10% and 7.19% in terms of *P_acc_* and *P_f_*, respectively. Comparing MsDANN and MsNN based EEG-AVE model performance under EEG+Visual and EEG+Audio embedding strategies, both *P_acc_* and *P_f_* values also have similar decrease patterns. The *P_acc_* value decreases from 74.65% to 70.04% for EEG+Visual, and from 69.08% to 65.33% for EEG+Audio. The *P_f_* value declines from 74.61% to 73.03% for EEG+Visual, and from 73.06% to 71.64% for EEG+Audio. When non-aggregated groundtruth is used, cross-comparing MsDANN and MsNN based model performance under an embedding strategy of EEG+Visual+Audio, it is found that the decoding performance significantly decreases from 71.13% to 63.58% (decreased by 10.61%) in terms of *P_acc_* and from 66.83% to 62.35% (decreased by 6.7%) in terms of *P_f_*. Similar trends are also observed in the other embedding strategies. The corresponding detection accuracies decrease to 60.98% (*P_acc_*) and 59.20% (*P_f_*) for EEG+Visual embedding strategy, and to 53.38% (*P_acc_*) and 56.34% (*P_f_*) for EEG+Audio embedding strategy. On the other hand, for arousal detection, the aggregated results show MsDANN based EEG-AVE model outperforms MsNN based EEG-AVE model across all three different embedding strategies in terms of both *P_acc_* and *P_f_*. For EEG+Visual+Audio, EEG+Visual and EEG+Audio embedding strategies, the corresponding improvement rates from MsNN to MsDANN are 9.18%, 7.30% and 4.93% for *P_acc_*, and that are 6.30%, 3.71% and 1.08% for *P_f_*. For non-aggregated results, similar patterns are observed. Better results are achieved when MsDANN based EEG-AVE model is adopted. Here, the improvement rates for three different embedding strategies are 14.60%, 14.73% and 10.73% for *P_acc_* and 8.73%, 7.90%, and 2.72% for *P_f_*.

Similar comparison results are also observed on the DEAP database, where MsDANN generally performs better than MsNN across all three embedding strategies (EEG+visual, EEG+Audio, EEG+Visual+Audio). For example, comparing MsDANN and MsNN based EEG-AVE model performance under EEG+Visual+Audio embedding strategy, the *P_acc_* and *P_f_* values of valence decrease from 75.26% and 77.16% to 66.03% and 72.55% for aggregated groundtruth and from 68.50% and 68.81% to 55.59% and 61.84% for non-aggregated groundtruth. The average decrease rate is 11.80%. For arousal, MsDANN outperformed MsNN when non-aggregated groundtruth is adopted.

The above results demonstrate that, comparing to MsNN, MsDANN is much more powerful in the proposed EEG-AVE model to deal with the problem of the individual differences in EEG signal processing. It provides a reliable and useful way to adaptively learn the shared emotion-related common and discriminant feature representation across individuals and demonstrates the validity of domain adaptation method in EEG-based affective detection applications.

### C. Performance Evaluation of Multimedia Representation

In this study, audio-visual information is represented by deep features characterized from two pretrained networks. We further verify the effectiveness of the deep feature representation and compare it with the performance using more traditional handcrafted features. Inspired from the previous video affective studies [22], [43]–[45], the commonly used handcrafted features are extracted and compared here. For visual information representation, the adopted handcrafted features include lighting key features, color information, and shadow portions in the HSL and HSV spaces. For audio information representation, the used traditional audio features include energy, loudness, spectrum flux, zero-crossing rate (ZCR), Mel-frequency cepstral coefficients (MFCCs), log energy, and the standard deviations of the above ZCR, MFCC, and log energy. The affective analysis of multimedia content with different feature representations is conducted and the corresponding comparison results of valence and arousal are summarized in Table V. The results show compared to the performance presented in Table IV, a significant improvement in affective detection performance is obtained when deep feature representation is used instead of handcrafted features. It reveals that compared to the traditional handcrafted feature representation, deep feature representation is a better and richer affective representation for understanding and perceiving the multimedia content.

**TABLE V:** Affective detection performance of MsDANN and MsNN on MAHNOB-HCI and DEAP databases using handcrafted features.

|  | Embedding Strategy | EEG Model | MAHNOB-HCI |  |  |  | DEAP |  |  |  |
| --- | --- | --- | --- | --- | --- | --- | --- | --- | --- | --- |
|  |  |  | Aggregated |  | Non-Aggregated |  | Aggregated |  | Non-Aggregated |  |
| | | | $P_{acc}$ | $P_f$ | $P_{acc}$ | $P_f$ | $P_{acc}$ | $P_f$ | $P_{acc}$ | $P_f$ |
| **Valence** | EEG+Visual | MsDANN | 62.68 | 64.75 | 51.38 | 46.23 | 59.96 | 58.80 | 61.40 | 57.04 |
|  |  | MsNN | 59.96 | 65.08 | 47.45 | 47.03 | 56.51 | 62.75 | 50.66 | 54.90 |
|  | EEG+Audio | MsDANN | 64.11 | 62.70 | 59.51 | 49.86 | 59.31 | 65.92 | 55.62 | 60.39 |
|  |  | MsNN | 61.82 | 64.42 | 54.41 | 50.37 | 56.52 | 66.89 | 48.74 | 58.65 |
|  | EEG+Visual+Audio | MsDANN | 73.52 | 69.26 | 64.99 | 50.48 | 61.05 | 63.65 | 59.89 | 59.82 |
|  |  | MsNN | 68.75 | 68.58 | 58.43 | 50.52 | 57.23 | 65.08 | 50.25 | 56.79 |
| **Arousal** | EEG+Visual | MsDANN | 70.04 | 71.99 | 55.23 | 50.87 | 64.56 | 76.64 | 46.73 | 59.27 |
|  |  | MsNN | 66.08 | 71.15 | 50.08 | 50.91 | 65.06 | 77.02 | 44.69 | 57.83 |
|  | EEG+Audio | MsDANN | 65.23 | 71.41 | 52.95 | 55.51 | 62.00 | 71.44 | 53.94 | 58.91 |
|  |  | MsNN | 62.51 | 70.97 | 48.03 | 54.11 | 62.99 | 72.36 | 48.77 | 54.64 |
|  | EEG+Visual+Audio | MsDANN | 72.18 | 72.92 | 59.79 | 53.71 | 61.69 | 67.74 | 58.93 | 58.00 |
|  |  | MsNN | 67.33 | 71.39 | 52.80 | 51.99 | 63.11 | 69.34 | 50.89 | 50.58 |

### D. Conclusion

In this paper, we propose a novel affective detection model (EEG-AVE) with an embedding protocol, where both EEG based emotional individual preferences and audio-visual based intrinsic emotions are incorporated to tackle the problem of the individual differences in EEG processing. The multimodal information is analyzed and compensated to realize efficient and effective EEG-based affective detection. The experimental results show that the proposed EEG-AVE model achieves promising affective detection results, comparing to the state-of-the-art methods. Besides, aiming at characterizing dynamic, informative, and domain-invariant EEG features across individuals, we develop a deep neural network with a transfer learning method (MsDANN) to solve the problem of the individual differences in the EEG data processing and investigate the performance variants with different neural network architectures (with or without domain adaptation). Our analysis demonstrates a superior cross-individual result is achieved under an evaluation of the leave-one-individual-out cross-validation individual-independent method. Furthermore, we utilize two well-known pretrained CNNs for semantic audio-visual feature extraction and introduce hypergraph theory to decode deep visual features, deep auditory features, and deep audio-visual fusion features for intrinsic emotions estimation. The possibility of affective detection using the multimedia materials is verified and the benefit of the proposed embedding strategy is examined. These results show both EEG signals and audio-visual information play important and helpful roles in affective detection, and the proposed EEG-AVE model could be applied to boost the development of affective brain-computer interface in real applications.

## VI. Acknowledgments

This work was supported in part by the National Natural Science Foundation of China under Grant 61906122, in part by Shenzhen-Hong Kong Institute of Brain Science-Shenzhen Fundamental Research Institutions (2021SHIBS0003), in part by the Tencent “Rhinoceros Birds”-Scientific Research Foundation for Young Teachers of Shenzhen University, and in part by the High Level University Construction under Grant 000002110133.

